# Exploring midwives’ understanding of respectful and non-abusive maternal care in Kumasi, Ghana: Qualitative Inquiry

**DOI:** 10.1101/708776

**Authors:** Dzomeku Veronica Millicent, Bonsu Adwoa Bemah, Nakua Kweku Emmanuel, Agbadi Pascal, Lori R. Jody, Donkor Peter

## Abstract

**Background:** Various aspects of disrespect and abusive maternity care have received scholarly attention because of frequent reports of the phenomenon in most healthcare facilities globally, especially in low- and middle-income countries. However, the perspectives of skilled providers on respectful maternal care have not been extensively studied. Midwives’ knowledge of respectful maternity care is critical in designing any interventive measures to address the menace of disrespect and abuse in maternity care. Therefore, the present study sought to explore the views of midwives on respectful maternity care at a Teaching Hospital in Kumasi, Ghana.

**Methods:** Phenomenological qualitative research design was employed in the study. Data were generated through individual in-depth interviews, which were audio-recorded and transcribed verbatim. Data saturation was reached with fifteen midwives. Open Code 4.03 was used to manage and analyse the data.

**Findings:** The midwives’ understanding of respectful maternity care was comprised of the following components: non-abusive care, consented care, confidential care, non-violation of childbearing women’s basic human rights, and non-discriminatory care. Probing questions to solicit midwives’ opinions on an evidenced-based component of respectful maternity care generated little information, suggesting that the midwives have a gap in knowledge regarding this component of respectful maternity care.

**Conclusion:** Midwives reported an understanding of most components of respectful maternity care, but their gap in knowledge on evidenced-based care requires policy attention and in-service training. To understand the extent to which this gap in knowledge can be generalized for midwives across Ghana to warrant a redesign of the national midwifery curriculum, the authors recommend a nationwide cross-sectional quantitative study.

## Background

The rise in facility-based deliveries with skilled providers in low-and-middle-income countries (LMICs) has resulted in decreased maternal and neonatal morbidities and mortalities [1–5]. In recent times, frequent accounts of facility-based disrespect and abusive care (D&AC) are undermining the purpose of encouraging childbearing women to access intrapartum care services in healthcare facilities [6–10]. D&AC violates childbearing women’s rights to quality maternity care, life, health, dignified care, and freedom from discrimination [11]. Preventing and eliminating facility-based D&AC in LMICs requires that countries implement a scalable, sustainable, and cost-effective solution. The World Health Organization (WHO) statement on addressing D&AC suggests that ensuring and integrating respectful maternity care (RMC) in obstetric care— pregnancy, childbirth, through postnatal care—is the appropriate solution to pursue [11]. RMC refers to “care organized for and provided to all women in a manner that maintains their dignity, privacy and confidentiality, ensures freedom from harm and mistreatment, and enables informed choice and continuous support during labour and childbirth” [12].

Studies have shown the effectiveness of RMC interventions in reducing D&AC and maternal and neonatal deaths in LMICs [13–15]. Interventions providing mentorship and training that transformed negative provider attitudes and improved the interpersonal communication skills of caregivers, along with interventions equipping facilities with efficient monitoring systems, and interventions educating childbearing women on their rights have had the most positive effect [14]. One intervention integrated specific components of RMC, like dignity, respect, communication, autonomy, and supportive care, into a simulation training [13]. The simulation training was designed to improve the identification and management of obstetric and neonatal emergencies [13]. It was piloted in the East Mamprusi District in the northern region of Ghana [13]. The findings showed that RMC training workshops have the potential to improve childbearing women’s childbirth experiences in LMICs [13].

To prevent and eliminate facility-based D&AC during childbirth, WHO recommended five key actions, one of which is to generate data related to respectful and disrespectful care practices [11]. In line with this, several studies have generated knowledge on the perspectives of childbearing women and skilled providers on D&AC [6,16–20]. The few studies that explored the views of skilled providers on RMC were from countries other than Ghana [21,22]. Thus, the present study seeks to explore the experiences and understanding of midwives on RMC in a tertiary health facility in Kumasi, Ghana. The aim of the study originates from the dearth of literature coupled with the evidence of incidences of D&AC from mothers in many healthcare facilities across the country [6,23–25]. Documenting the thoughts of midwives on RMC will help policymakers to address existing gaps in knowledge on RMC. It will also provide further evidence of non-evidenced based practices in maternity care.

## Design

Qualitative research design with a phenomenological approach was employed to explore the experiences and views of midwives about respectful maternity care. The phenomenological approach was used because it permits the authors to explore and document midwives’ views of RMC [26]. This study is part of a larger qualitative research study exploring the feasibility of changing the culture of disrespect and abuse in maternity care.

### Study Setting

The study setting was a tertiary health facility in Kumasi, located in the Ashanti region, the central part of Ghana. The study setting was chosen because of the major role it serves in Ghana’s healthcare delivery. The tertiary health facility in focus serves patients across the country and has a bed capacity of approximately 1,200 and a staff strength of about 3,000. It is the main referral hospital for six political and administrative regions in Ghana—the Ashanti, Brong Ahafo, Western, the three Northern regions (Northern, Upper East, And Upper West)—, and neighbouring countries. It has twelve (12) clinical directorates, one of which is the maternal and child health directorate (MCH-D). The MCH-D has two main units—the general ward and the special ward. The general ward is an open ward the use of which is covered under national health insurance, and the special ward is a private ward which is accessed at an extra cost.

### Population, Sampling, and Sample Size

The study population was composed of midwives in this tertiary health facility in Kumasi, Ghana. Inclusion criteria were midwives working on the labour wards for at least one year. Purposive sampling ensured that participants with expertise were enrolled. The first author and two research assistants (RAs) recruited the study participants. All participants who officially agreed to be included completed the study, and saturation of data was achieved with fifteen (15) midwives, at which point no new themes or sub-themes emerged [27,28].

### Data Collection

A semi-structured interview guide and face-to-face in-depth interviews were used to generate the data. The researcher and the RAs asked probing questions to acquire an exhaustive and in-depth understanding of the participant views regarding RMC. Data collection started on 3^rd^ January 2019 and ended on 25^th^ February 2019. The interview guide was pre-tested with 3 midwives working at the maternal unit of the Kwame Nkrumah University of Science and Technology Hospital, Kumasi to ensure the appropriateness of the guiding questions. The interviews were conducted in English, but participants were permitted to express themselves in Twi, the native language of the region. All data collectors spoke English and Twi fluently. The interview duration was 50 - 80 minutes, and the interviews were audio-recorded with the participants’ consent. Date, time and venue of the interviews were scheduled to be suitable for participants. During the interview sessions, the researcher took field notes of non-verbal cues and other relevant observations.

### Data management and Analyses

The researcher and the RAs thoroughly listened to the audio files for accurate verbatim transcription. The transcribed interviews were proofread to ensure that participant’s views were precisely captured. Anonymity was ensured by serializing each transcript file, and the transcripts were kept in a secured folder on the laptop of the principal investigator. Open Code 4.03, a qualitative data management software, was used to manage and analyze the data by adopting thematic analysis principles. The D&AC project was created in the software, and the transcripts were saved as text files and imported into the project folder. Each transcript was coded, and the codes synthesized into subthemes and further into themes base on their similarities and relationships [29,30]. The emergent themes structured the presentation of the findings.

### Trustworthiness/Rigour

Confirmability, transferability, dependability, and authenticity were the trustworthiness criteria followed in the study [31]. Adopting purposive sampling techniques ensured that participants who had relevant midwifery experience were recruited. Confirmability was achieved through member checking to help ensure participants’ realities were accurately presented. Also, independent analysis of the data by the authors further confirmed the findings. To ensure transferability, the authors provided a detailed description of the research design and data collection methods and setting as well as the background of the participants. Through peer debriefing and strict adherence to the study protocol, the trustworthiness of the data was further ensured. Member checking was done to confirm that participants realities were truly presented before drawing the final conclusions of the data [31].

### Ethics approval and consent to participate

The Committee on Human Research, Publication, and Ethics (CHRPE) at the Kwame Nkrumah University of Science and Technology (KNUST) gave the ethical clearance for the study (CHRPE/AP/181/18), and the managerial board of the teaching hospital gave the institutional approval for the study (RD/CR17/289). Participants were briefed on the study and their rights to voluntary participation and withdrawal from the study with no consequences. Only participants who consented were involved in the study. The participants gave their consent to the interview, the audio recording of the session, and the publication of findings. Participants’ confidentiality was ensured by conducting the interview in an enclosed office. Information that could reveal the identities of the participants were excluded from the transcripts to ensure participants’ anonymity.

## Findings

### Demographic Features of Participants

The midwives were, on average, 33 years old, with a range of 31-48 years. They had engaged in professional practice for an average of eight years. Seven participants obtained a bachelor’s degree in midwifery, while nine participants received a diploma [diploma is a post-secondary qualification that’s lower than a bachelor’s degree in Ghana’s educational system]. Only one of the midwives was a Muslim, and the others were Christians. Eleven were currently married. Those with children (n=10) had an average of 2.3 living children (range = 1-3).

### Respectful and Non-Abusive Maternity Care (RMC)

Views of the midwives were sought on respectful and non-abusive maternity care (RMC). The following themes emerged from the analyses: awareness of respectful and non-abusive maternity care, motivation for RMC, and recommendations for optimal RMC. The sub-themes associated with the themes are reported in table 1.

**Table 1:** Themes and Subthemes

| Themes | Subthemes |
| --- | --- |
| **Awareness of RMC** | 1. Respectful care |
|  | 2. Non-abusive care |
|  | 3. Provision of childbearing women-centred care |
|  | 4. Respecting childbearing women's rights |
|  | 5. Evidence-based care |
|  | 6. Illustrations of RMC |
| **Motivation for RMC** | 1. Childbearing women's wellbeing |
|  | 2. God's calling |
|  | 3. Economic, social, and psychological benefits |
|  | 4. Midwives' experience of labour |
| **Recommendation for Optimal RMC** | 1. Labour and pain management education |

### Awareness of RMC

All the midwives in the current study demonstrated some understanding of respectful and non-abusive maternal care. The sub-themes associated with the description of RMC include respectful care, non-abusive care, ensuring childbearing women-centred care, respecting childbearing women’s rights, and illustrations of RMC.

#### Respectful care

The midwives have knowledge of comfort measures that the childbearing women may require to cope with the process of labour. Some of these measures include consent seeking, clinical procedure explanation, and providing sacral massage. These are illustrated in the following responses of midwives:

*You [the midwife] are there to support that childbearing woman at that moment, so, you assist in sacral massage, and you give her encouraging words in order that the childbearing woman will not feel that she is in that alone…there are a lot of childbearing women in there, so we are in as their support to go through their labour and the pain…So, we are the caregivers and we are also the supporters …we are giving them the support during the labour.* [Midwife 001].

*You must show love and welcome her, so she feels she has come home. In that way, even if she came with some anxiety, she will become relaxed…I consider it [consent seeking] as respect because if you are just there, and out of nowhere, someone tells you to turn to get an injection, even if you are not afraid, you could be scared a little. But if a worker comes and explains a procedure to you, for instance, because of the headache you are experiencing, this injection will take absolute care of it, and then she seeks your consent to give you the injection, it shows respect. …* [Midwife 004].

### Non-abusive care

The midwives described and gave examples of non-abusive care that encapsulated providing the care that does not compromise the physical, psychological dignity, and wellbeing of childbearing women. The key phrases in their descriptions include reassuring childbearing women, ensuring childbearing women’s comfort, desisting from the use of harsh or abusive language, and the use of non-verbal communication. Their views are presented as follows:

*Non-abusive [care] is rendering care that will not make the* childbearing woman *feel bad. Yeah, care that will make the childbearing woman feel comfortable at your ward. Yeah. You ensure by the way you talk to the childbearing woman nicely and encouraging, reassuring, or make the childbearing woman comfortable.* [Midwife 007].

*Non-abusive care is the care you give to childbearing women without using abusive words, like insults or shouting or using non-verbal languages to communicate with childbearing women.* [Midwife 013].

### Childbearing women-centred care

In describing RMC, most of the midwives mentioned and offered practical examples of individual-or childbearing woman-centred care. It involves taking into consideration the childbearing women’s unique experiences, feelings, and response to labour and care.

*Respectful care is taking the woman who comes into labour, the feelings into consideration so that no matter what she does, you will respect her and do the best for her.* [Midwife 002].

*Well, there are a lot of clients who come here. And each one has peculiar characteristics. Let’s use labour as an example. Someone will go into labour quietly. If you don’t get close to her, by the time you realize, she will deliver the baby without you. But someone else will shout and wail from 2cm till the end. So, each person is different. As a result, you don’t claim someone came and was quiet so another person should be quiet; you should be able to care for each childbearing woman devoid of discrimination. You first determine the peculiarities, and then give her appropriate care that will help make the childbearing woman better.* [Midwife 004].

### Respecting childbearing women’s rights

Knowing that respecting childbearing women’s rights is a relevant component of RMC, the researchers explored the awareness of midwives on the rights of childbearing women during care. The analyses revealed that the midwives were aware of the rights of childbearing women during care. They indicated that childbearing women, first and foremost, have the right to RMC. Other rights to which midwives believed childbearing women were entitled include the right to confidentiality and anonymity of health status, the right to consented care, the right to have access to relatives during the stay at the hospital, the right to health information, the right to privacy, and the right to choose from treatment options. The following are some of the views of the midwives on childbearing women’s rights:

*The right to care, the right to information, the right to privacy, the right to the best of care and the right to consent to any procedure supposed to be performed on her, and the right to get an explanation concerning any procedure supposed to be performed on her …Ideally, before you administer any drug to a childbearing woman, the first thing you do is to explain what the drug is going to do and then the consequences attached to it.* [Midwife 002].

*…A single room like here [pointing to a room], and she wants a relative to visit or be around her, suppose she’s never slept alone in a room before, she has the right to have a relative be around as long as there is room for that to happen…Also, there are certain kinds of information, though she may have come with her husband, she has a right to demand that certain information remains between you and her; her husband doesn’t even have to know. She also has that right…And then, for instance, a childbearing woman can ask for an alternate, say an oral medication instead of an injection because she may not like injections. For that too, she has a right because she can’t take an injection and as long as an alternative exists that she can orally ingest, she has that right. Also, any information concerning her care, she has every right to have access to it…* [Midwife 004].

### Evidence-based care

The views of midwives were sought on evidence-based care, but their inability to respond to the questions triggered further probing. The midwives who responded to the probing questions mentioned that hitting the thighs of the childbearing women in the second stage of labour helps them push and deliver babies safely. Their views on hitting childbearing women during labour carried moral overtones, as they judged the action to either be wrong or right. Their concerns were not about whether the practice was evidence-based. In connection with evidence-based care, the interviewer asked one of the midwives to share her opinion on hitting childbearing women. This is what the participant had to say regarding hitting a childbearing woman:

*when the head is crowned, then you have to hit the childbearing woman [in labour]. If you don’t, she will make you lose the baby...and people will blame you or call you names. Before that, you have to advise the childbearing woman to push. But there are some that you will advise, but they will not take it. But once you hold the thigh like this [demonstrating pinching], they become conscious. And when they are done, they even apologize to you...at that particular time, if any midwife does this, she hasn’t done anything wrong, because at that time, either we save the mother, or we save the baby*.[Midwife 008B]

Some midwives were not sure whether hitting childbearing women was evidence-based, but they felt such a practice was not right.

*...Well, it’s bad to hit childbearing women…I...only [do it] when conducting delivery and she is closing the gap and I hit in-between the thighs ‘open up!’, aha, that’s the only time I hit a childbearing woman, and it is not hitting deliberately…*[Midwife 003]

*...I’ve not done some before but I think it’s not good to hit a childbearing woman.* [Midwife 008]

### Illustrations of RMC

The midwives presented some examples of their personal experiences of providing RMC. From their testimonies, it is evident that some of the midwives empathized with the childbearing women, especially those who exhibited unexpected behaviours, and this resulted in the provision of RMC. Some also went out of their way to provide relief for economically disadvantaged childbearing women. Instances where childbearing women were unable to pay their bills, the midwives either, made the necessary provision from their pockets or connected them with the district’s social welfare department for assistance. These are captured in the following quotes:

*We sometimes made solicit for funds to buy malt, cord sheets and such for some [poor] childbearing women. The belief is that God even blesses you when you treat such less-endowed individuals nicely, so you have no reason to insult such people and treat them badly because they have no money; you just deliver your God-ordained duty. I was always soliciting for things for such people and some of my colleagues would make fun of me going around begging for items, but it was nothing. A childbearing woman has given birth and may collapse from hunger, so going around asking for help to feed her properly was no big deal.* [Midwife 004].

*Now, the hospital policy is that, the staff of social welfare should intervene and help the childbearing woman out.* [Midwife 005].

*Sometimes we do help them. Maybe if you have money with you, you do buy some of the cord sheets for the person, the items for the person. And if the relatives come, and they can afford to pay you back, then you take your money but if they can’t, you just let it go.* [Midwife 008].

Some midwives cited examples of the positive outcome of involving relatives of childbearing women in the caregiving process. From the examples, it’s clear that relatives’ involvement had made easy the provision of optimal maternal care.

*Most of the time, they come with their relative to support them but because it is an open ward, we ask the relatives to visit their family member who’s in our care. We have a particular time they have to come in; that’s when the visiting times are.* [Midwife 001].

*Anytime they wanted to perform any procedure, if she tells the husband that they want to take x-rays and that stuff, he would not agree; he would say he doesn’t have money. So, I had to pick the guy’s number, call him, let him come to the ward, we had a nice chat. So anytime they wanted to do anything for her, I’ll just call the husband, ‘today, we are doing this procedure for her,’ and within one (1) hour or two (2) hours, he will bring the money. Sometimes, he will even send me money through mobile money, [and I will] pick it and go do the procedure for her.* [Midwife 002].

### Motivations for RMC

The midwives had certain motivations for providing childbearing women with RMC. Their motivations centred on the childbearing women’s wellbeing, the midwifery profession as God’s calling, the economic, social, and psychological benefits associated with RMC, and the experience of labour.

#### Childbearing women’s wellbeing

The midwives interviewed had experiences that compromised or enhanced the wellbeing of childbearing women. Thus, they understood childbearing women’s reactions to both abusive and non-abusive care. From the midwives’ views, it’s revealed that childbearing women’s wellbeing is one notable motivation for providing RMC. According to the midwives, childbearing women remain appreciative to them due to the RMC they received.

*The most important thing is to deliver a healthy baby, but that ultimately depends on the kind of care rendered to the mother. So, if the childbearing woman is wet, she must be wiped and changed. …The baby inside the mother also experiences whatever you do for the mother. The baby needs to enter this world with joy and that depends on the conditions the mother experiences. So, if you educate her to do everything correctly, the pushing becomes exceptional when the time is due; the mother has the energy to push and the baby is strong.* [Midwife 004].

#### God’s calling

Some of the midwives noted that God called them to serve as midwives, and that motivates them to provide optimal RMC. They had the following to say:

*I believe God has given me this opportunity [serving as a midwife] so I won’t let Him regret giving me the blessing.* [Midwife 004].

*What I want to share with them [colleague midwives] is that this job is a calling, it’s from God and if you do it well, the blessing is even here on earth. And some people say when you do well here, when you get to heaven you will get it back, but we get the reward, whether good or bad, we get them here.* [Midwife 005].

#### Economic, social, and psychological benefits

The midwives indicated that the benefits they have reaped due to providing RMC encouraged them to continually provide RMC. The benefits were economic, social, and psychological in nature. Paramount among the benefits they accrued from providing RMC was emotional appraisal from postnatal women and their relatives.

*They [the childbearing women] really remember whatever you do to them. Sometimes you meet them in town, and they call you and you feel happy. They sometimes say, “This is your grandchild,” and you are like, wow! “I haven’t even given birth yet, but I have a grandchild.” It’s beautiful when they call you. Sometimes, call you to find out how you are doing… [Sometimes, they will say] “Oh bring your clothes and let me sew them for you.”* [Midwife 001].

*…If a childbearing woman is grateful for some work you do for her, and she is showing appreciation when it is time to be discharged, that is wonderful…Right there, and I can get very happy. In some of such encounters, I may be walking with my husband and would commend me saying ‘you see the work you’ve been doing is bearing fruits’ and I will respond by ‘yeah, that’s what protects us’…* [Midwife 008B].

*Sometimes when the client comes, in my experience when they come, and you are free with them, you talk to them very nicely, even when they have given birth, they are coming for review, they see you, they just approach very nicely and they ‘Oh, God bless you. This midwife, when I came to deliver, she was exceptionally good to me. This midwife has a welcoming demeanour’. It is very good when people use such words to describe you. Even the relatives, when they come, they talk to us and we receive them properly.* [Midwife 009].

#### Midwives’ experience of labour

One midwife, who had an experience of giving birth, noted that experiencing labour puts you in a position to provide RMC. This is what she said:

*I once heard that if you’d never given birth before, you were never allowed into midwifery. But there came a time when they needed more people and all these young people were allowed in. Because if you’ve ever had babies [you will appreciate the need to provide good care to childbearing women going through similar labour experience] …when I was in the labour ward, my colleagues always complained that I had too much tolerance for bad behaviour.* [Midwife 004].

### Recommendation for optimal RMC

Recommendation for optimal RMC was the final theme, and labour and pain management education as the associated subtheme.

### Labour and pain management education

Some of our participants suggested that, to ensure optimal RMC, during antenatal care visits, childbearing women should be thoroughly equipped with information on the phases of labour and pain management. They believe that this education will help childbearing women to cope with and manage their pains during labour and make them cooperative with caregivers when admitted at the facility. The following are the suggestions from the midwives:

*…when they are pregnant, we should give them every education they need to know, every information they need to know. Sometimes, they don’t even know how to even attach the baby to the breast, which is very bad. But if from antenatal she is knowing that this is how labour is, this is how when the baby is about coming, this is how it is, this is how you have to attach your baby to the breast, “even we ourselves, we won’t get tired/frustrated” because she knows so she will act according to that information she is having….* [Midwife 001].

*Sometimes, they need to be talked to, especially during the antenatal. We need to educate them on what they are supposed to do when they are in labour. If we do that very well, when they come to labour, they will know what they are supposed to do…the person [childbearing woman] comes to labour not knowing anything about the first stage, second stage or third stage. All she knows it; she has come to give birth. We have to educate her, so she understands what she is coming to accomplish. So next time she is coming, she knows this sister said when I do get here, I will do this, I will do this. So, don’t say ‘she came to deliver, and it is done’, no. sometimes you have to educate her on the baby.* [Midwife 008].

## Discussions

This study explored the views of midwives on RMC. Findings indicated that midwives at the study site understood the following components of RMC: non-abusive care, consented care, confidential care, non-violation of childbearing women’s basic human rights, and non-discriminatory care. The study highlighted a gap in the knowledge of RMC on evidence-based care.

Midwives in this study have substantial knowledge of respectful and maternal care, but their support for non-evidence-based practices suggests that knowledge on RMC may not always translate into delivery of quality maternity care. Other studies in healthcare facilities in Ghana have revealed that childbearing women are usually disrespected and abused, and the midwives in our study and in others justified D&AC by arguing that childbearing women should not be overly pampered as this may lead to non-compliance and death of their babies [6,23,25].

This finding from our study further suggests that equipping midwives with knowledge on RMC alone without identifying and addressing other potential facilitators of D&AC cannot eliminate and prevent D&AC. Evidence suggests factors that might promote RMC in LMICs include creating access to quality training and supervision for midwives, increasing staff numbers and reducing workloads, improving salaries and living conditions of midwives, building well-equipped, well-organised healthcare facilities with all the essential resources needed for quality care, fostering teamwork, trust, collaboration, and communication among health workers and with childbearing women [32,33].

The midwives made reference to intrinsic and extrinsic motivations for providing RMC. These motivations include an emotional appraisal, gifts from postpartum women, and the regarding of midwifery profession as God’s calling. This finding is congruent with findings obtained from studies in Ghana, Burkina Faso, and Tanzania [34,35]. These studies indicated that midwives were motivated to provide RMC because of appreciations received from the community, positive feedback from superiors, perceived government and NGO support, and their sense of duty towards childbearing women and God [34,35].

Midwives also mentioned that educating childbearing women about labour and pain management will ensure optimal RMC. This recommendation is an example of how quick midwives are to outline measures directed at childbearing women to curtail D&AC. It’s laudable to provide quality pain management education to childbearing women during antenatal care visits; however, the training of midwives to learn to provide childbearing women with comfort measures that help them cope with and manage pain during the experiences of childbirth is an essential strategy for promoting respectful maternity care [13].

### Limitation and strength

One obvious limitation of this study is the issue of generalizability of findings for other midwives in the teaching hospital itself, the region of the study setting, and the entire country. This limitation stems from the exploratory nature of the research design. Another limitation is that the midwives may have been biased either about their own abilities and use of RMC or about how childbearing women respond to it. However, a significant strength of this study is the obtaining of comprehensive information on midwives’ perspectives on RMC and the identification of skilled providers’ gap in knowledge on evidence-based maternity care. Another important strength of this study is the richness of the data; maternity care providers—midwives—gave their opinions on how they think and implement RMC care, which is critical in both understanding and improving this care.

## Conclusions

The objective of the study was to explore midwives’ experiences of and views on respectful maternal care. The findings revealed that the midwives have an understanding of most components of respectful maternal care, but their gap in knowledge on evidenced-based care requires policy attention and in-service training. To understand the extent to which this gap in knowledge can be generalized for midwives across Ghana to warrant a redesign of the national midwifery curriculum, the authors recommend a nationwide cross-sectional quantitative study.

## Acknowledgement

We acknowledge the midwives who participated and shared their views and experiences on RMC. Pre-Publication Support Service (PREPSS) supported the development of this manuscript by providing pre-publication peer-review and copy editing.

